# Insulin-like growth factor 1 is related to the expression of plumage traits in a passerine species

**DOI:** 10.1101/645226

**Authors:** Katharina Mahr, Orsolya Vincze, Zsófia Tóth, Herbert Hoi, Ádám Z. Lendvai

## Abstract

Avian plumage colors and ornaments are ideal models to study the endocrine mechanisms linking sexually selected traits and individual parameters of quality and condition. The insulin-like growth factor 1 (IGF-1), an evolutionarily highly conserved peptide hormone, represents a link between body condition and the individual capacity to grow elaborated ornamental features, due to its regulatory role in cell proliferation and differentiation and its high sensitivity to the nutritional state of individuals. We investigated whether IGF-1 levels during molting affect the expression of multiple ornaments in a sexually dichromatic passerine species, the bearded reedling (*Panurus biarmicus*). We collected blood samples of males and females shortly before the molting completed and measured the size and colors of ornamental traits. Our results indicate that in males, structural plumage colors, the size of the melanin based ornament (beard) and tail length are independent traits. IGF-1 levels predict the length of the tail and the expression of male structural plumage components (UV-coloration), but not the melanin based ornament. In females plumage color and tail length were independent traits, which were not related to IGF-1 levels. Overall, our results indicate for the first time that IGF-1 could play a role in the development of secondary sexual characters in a passerine species.

## Introduction

The physiological processes shaping the relationships between showy ornaments and individual parameters of quality and condition have been investigated intensively over the past decades. In that respect, hormones and their pleiotropic effects present one of the key mechanisms, linking individual physiology and the capacity to develop elaborated sexually selected traits (Flatt & Heyland, 2011, Hudson & Wilcoxen, 2018, Laucht & Dale, 2012, Nolan Jr. et al., 1992, Peters et al., 2006, Roberts et al., 2009). The best studied examples are androgens, which orchestrate the investment into behaviors related to reproduction and stimulate the growth of secondary sexual characters (Alonso-Alvarez, 2001, Andrew, 1969, Flatt & Heyland, 2011, Peters et al., 2006, Roberts et al., 2009). Their effects, however, come along with costs, which result in suppression of the immune system and an increase in oxidative stress, which force the bearer of an ornament into a trade-off between the differential allocation of resources into ornamental features or self-maintenance (e.g. growth, immune-competence) (Flatt & Heyland, 2011, Hau, 2007, Ketterson & Nolan, 1999, Nolan Jr. et al., 1992). The capacity of individuals to cope with these trade-offs might constitute one of the key factors linking condition and ornament expression in many vertebrate species.

Although much less investigated than the sex steroids, another, phylogenetically more ancient hormonal pathway, the insulin/insulin-like growth factor 1 (IGF-1), plays a crucial role in the mediation of such life history trade-offs (Barbieri et al., 2003, Dantzer & Swanson, 2012, Harshman & Zera, 2007, Shit et al., 2014, Sparkman et al., 2009, Zera & Harshman, 2001). IGF-1, an evolutionarily conserved polypeptide metabolic hormone regulates growth, development and reproduction in vertebrates by stimulating cell proliferation, migration, differentiation, and protein synthesis (Dantzer & Swanson, 2012, Doublier et al., 2000, Liu et al., 1993). External factors such as stress, infection and nutritional status affect IGF-1 secretion and its effects on individual physiology (Dantzer & Swanson, 2012, Emlen et al., 2012, Lodjak et al., 2016, Tóth et al., 2018). Because of this sensitivity and its regulatory functions, IGF-1 plays an important role in shaping life history traits, which also involve the development of sexually selected characters (Ditchkoff et al., 2001, Emlen et al., 2012, Lewin et al., 2017, Suttie et al., 1985).

We propose that these properties make IGF-1 an interesting candidate to link parameters of body-condition and the expression of avian plumage ornaments. Ornamental plumage traits appear in different forms (Andersson, 1994b, Andersson et al., 2002, Hill & McGraw, 2006a, Hill & McGraw, 2006b, LaFountain et al., 2015, Roulin, 2016) and, despite their remarkable diversity, they share the same characteristics: they serve as “quality-indicators” to conspecifics and hence significantly affect individual access to resources and future reproductive success (Andersson, 1994a, Delhey & Kempenaers, 2006, Hill & McGraw, 2006a, Hill & McGraw, 2006b, Hudson & Wilcoxen, 2018, Jacot & Kempenaers, 2006, McGlothlin et al., 2007, McGraw et al., 2002, Murphy & Pham, 2012, Musgrove & Wiebe, 2016). Intensity, size and elaboration of plumage colors and ornaments are determined during molt and most types of ornamentation are costly to produce and therefore are highly sensitive towards parameters of condition during molting (Griggio et al., 2009, Hudson & Wilcoxen, 2018, Loyau et al., 2005, McGraw et al., 2002, Svensson & Merilä, 1996). The renewal and growth of feathers, is a long and energy-demanding process, which requires major changes in the metabolic rate, a vast increase in cell proliferation rate, cell differentiation and body protein synthesis (Kuenzel, 2003). Considering that IGF-1 (i) regulates these processes, which are tightly linked to the physiological requirements of molting, (ii) is highly sensitive towards nutritional status (Clemmons, 2012, Dantzer & Swanson, 2012, Gunnell et al., 2003, Mazzuco et al., 2005) and (iii) signals availability of resources (Bartke et al., 2003, Mattson et al., 2004), it might also affect individual capacity to grow condition dependent ornamental features. Whereas the role of IGF-1 in growth and development of avian species has been investigated (Beccavin et al., 2001, Lodjak et al., 2017, Lodjak et al., 2014, Lodjak et al., 2016, McMurtry et al., 1997), to the best of our knowledge, the relationship between plumage ornaments, body condition and IGF-1 concentrations during the molt has yet not been studied.

Conspecifics, often evaluate individuals based on several cues and signals, rather than one single trait, and in many species, males and females display multiple ornaments (Alonso et al., 2005, Andersson, 1994a, Andersson et al., 2002, Candolin, 2003, Mahr et al., 2016). These can provide redundant information and/or amplify a signal, provide different information or have no further informational content (“unreliable” signal) (Andersson et al., 2002, Candolin, 2003, Griggio et al., 2016, Ornelas et al., 2009). Furthermore, individuals of many species often carry ornaments and colors of different physiological origins, which might affect the information these traits convey to receivers (Hill & McGraw, 2006a). Therefore, in order to gain insight into the role of IGF-1 as possible link between parameters of condition and ornament expressions, we investigated multiple ornamental plumage traits. Our model species is a free-living European passerine, the bearded reedling (*Panurus biarmicus*). While displaying a strong sexual dichromatism, males and females carry multiple ornamental features, with different physiological origins (Figure 1). Male bearded reedlings are characterized by a distinct melanin based ornament, the black beard, which is an honest signal and underlies inter- and intrasexual selection processes. Previous studies on this species also demonstrated that the length of the tail is a sexually selected trait in males and females (Griggio et al., 2016, Hoi & Griggio, 2008, Peiró et al., 2006, Romero-Pujante et al., 2002). Males are also characterized by a conspicuous blue head, a rose/pink flank region and, both males and females, possess an achromatic bright chin (Figure 1).

**Figure 1.**
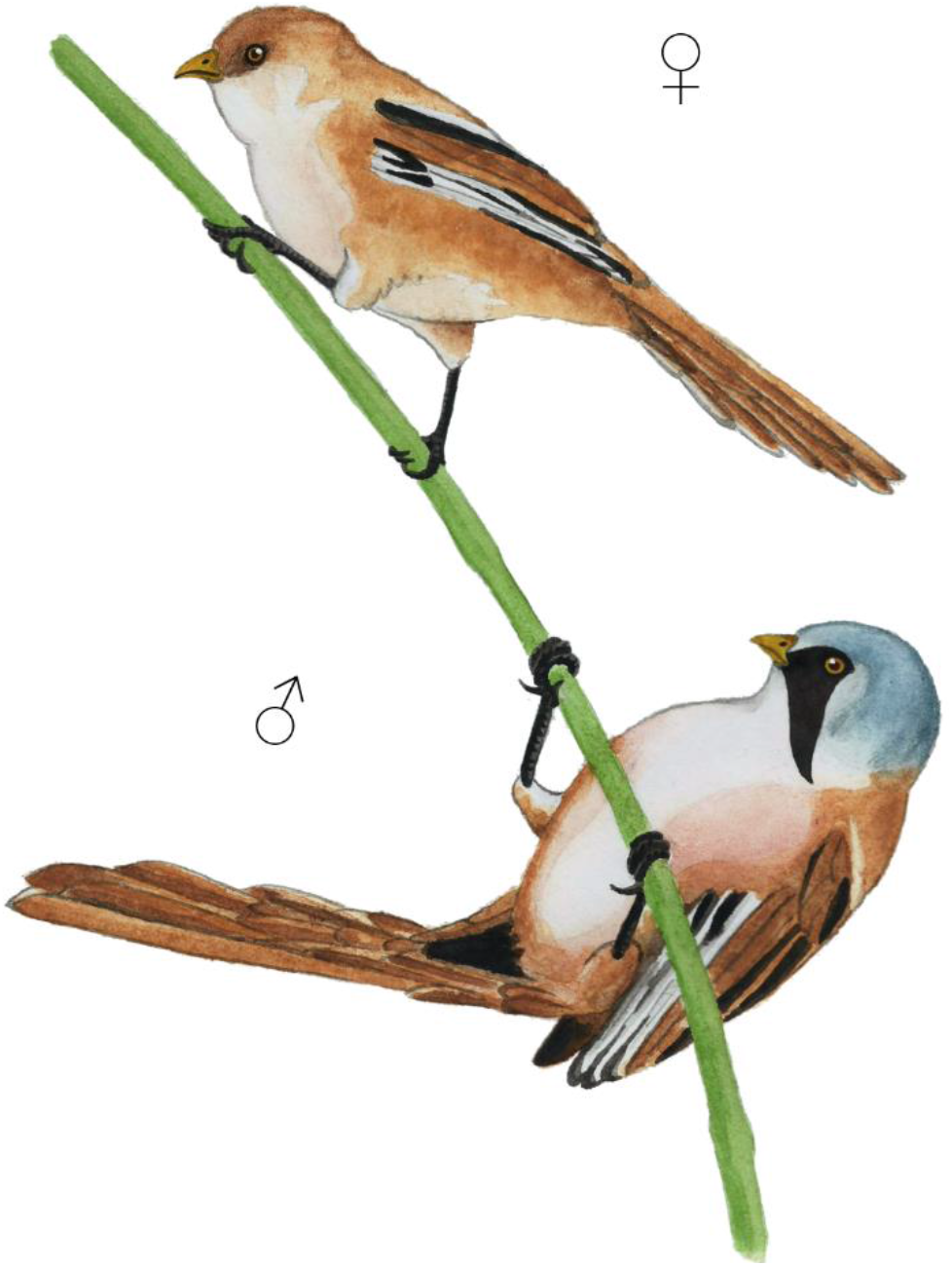
Female and male bearded reedlings display a strong sexual dichromatism. The conspicuous blue head and rose flank region of the males as well as the achromatic bright chin reflect in the UV-range (Original Artwork provided by G. Rédai, Department of Evolutionary Zoology and Human Biology, University of Debrecen, Debrecen, Hungary).

These plumage regions are characterized by reflection in the UV-range (Figure 2), which indicates the presence of structural plumage components. Structural plumage colors are the result of keratin structures and air-spaces embedded in the spongy medullary layer of the feather. Their regularity, which is highly sensitive towards parameters of individual condition during the molt predicts the degree of UV reflectance in the feathers (Griggio et al., 2009, Griggio et al., 2010b, Keyser & Hill, 1999). We therefore examined structural plumage components, by measuring UV-Chroma in addition to the melanin based plumage colors of the body feathers in both sexes.

**Figure 2.**
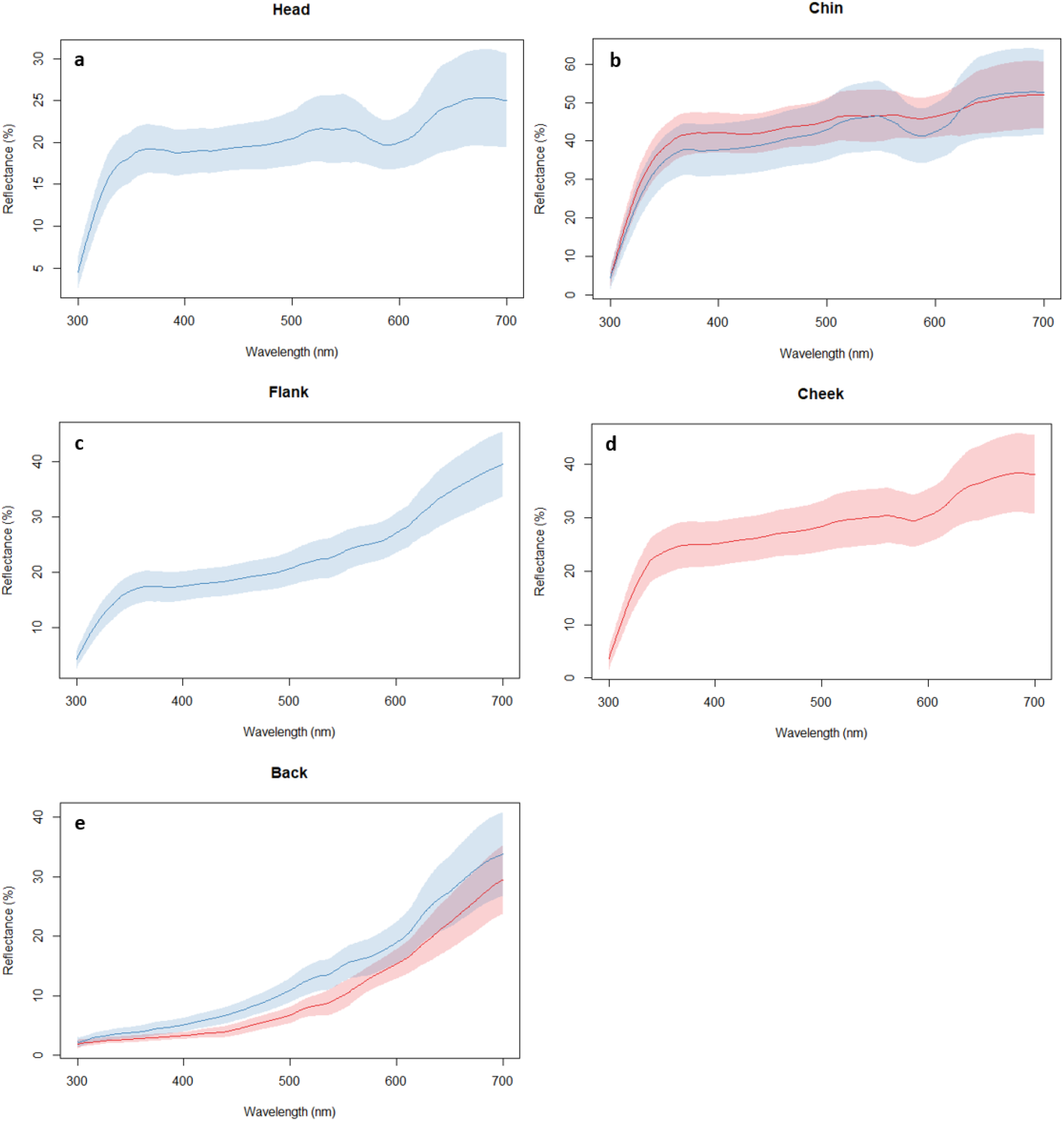
The average spectral curves based on 23 male (blue) and 20 female (red) bearded reedlings. Note that the head, flank, chin and the cheek (panels a-d) indicate reflectance in the UV-range (320-400 nm), while the brown coloration of the back (panel e) does not reflect in the UV range. The lines show the average reflectance and the shaded areas show the corresponding 95% confidence intervals.

In order to explore whether IGF-1 serves as possible link between body condition and plumage traits, we captured free-living bearded reedlings at the final stage of their molt and measured their baseline IGF-1 levels. Then we brought them into a spacious aviary under semi-natural conditions and after completion of the molt, we measured plumage traits in both sexes. We then tested for a relationship between the multiple ornamental traits and parameters of body size and condition (Griggio et al., 2016, Hoi & Griggio, 2008) and investigated whether baseline IGF-1 levels collected during late molting stages predicted the expression of plumage characteristics. We expected that ornamental features, which are sensitive towards body condition during molt would be related to condition parameters. If IGF-1 increases individual capacity to invest into the development of sexually selected characters, we expected to observe a positive relationship between IGF-1 plasma concentrations and parameters of plumage quality.

## Methods

### General methods

Free-living bearded reedlings were captured at the Lake Neusiedl (47°46’10.5”N, 16°45’20.1”E, Burgenland, Austria) in September and October 2016, by conducting mist-netting with between 0700-1600. Immediately after the capture, we collected blood samples (70-140 μl) by puncturing the brachial vein. The blood samples were drawn within 3 minutes (average time: 2 min 19 s) from the time when the bird hit the net, and therefore these samples are considered as baseline measurements (Romero & Reed, 2005, Tóth et al., 2018, Wingfield & Romero, 2001). Immediately after sampling, we transferred the blood into 0.5 ml microcentrifuge tubes, which were stored in a cooling box until further processing in the laboratory. The plasma was separated from red blood cells, by centrifuging the sample for 5 minutes at 2000g, and removing the plasma with a Hamilton syringe. We stored the samples in a −20°C freezer until assayed for IGF-1 by enzyme linked immunosorbent assay (ELISA; see details below).

All birds were banded with a unique combination of darvic color rings. Only individuals, which have molted more than two thirds of their plumage, entered the study; the sex was determined by plumage coloration (Svensson, 1992). Subsequently, we measured body mass (to the nearest 0.01 g), tarsus length (to the nearest 0.01 mm) and tail length (to the nearest 0.5 mm). Immediately after the measurements, we transferred the birds to individual cages and recorded their feeding behavior, by counting the amount of mealworms consumed within 2 hours. Only feeding birds were regarded as capable to cope with the captive situation and were transferred to the housing facilities of the Konrad Lorenz Institute of Ethology, University of Veterinary Medicine in Vienna, Austria. The birds were kept in a mixed sex flock in a large outdoor aviary (10 × 5 × 4 m), under natural light regime and conditions simulating their natural environment. The aviary contained dense vegetation and the birds received water, a mixed diet of commercial insectivorous food (protein mash, apples, quark, egg, carrots), seeds (canary seeds, millet, hemp seeds) and mealworms *ad libitum*.

### Characterization of plumage traits

We characterized plumage traits in male and female bearded reedlings after all individuals had finished molting. The freshly molted plumage coloration was measured using a USB-2000 spectrometer and a DHS-2000-FHS deuterium halogen lamp, connected through a bifurcated fiber-optic probe (Ocean Optics, Eerbek, The Netherlands). By fitting a black rubber cylinder on the top of the probe, we minimized disturbance by outer light sources and ensured a standardized distance and angle (90°). Prior to each measurement, the spectrophotometer was recalibrated. For the calibration of white we used a white standard (Avantes, Eerbek, Netherlands); for black, we removed the probe from the light source and closed the cap of the plug (Griggio et al., 2009, Mahr et al., 2016, Mahr et al., 2012).

We obtained 3 measurements from distinct regions of the plumage in males (head, flank, back, chin) and females (chin, cheek, back) (Figure 1). Based on the obtained spectral curves (Figure 2) and the literature, we focused on two standard descriptors of reflectance variables, which were generated from the raw reflectance data: brightness and UV-Chroma. We used brightness to determine the degree of coloration of the brown back of males and females (Figure 2), which we calculated as the average percent reflectance in the 320–700 nm range. Brightness was previously shown to reflect the melanin content of plumage features, with low brightness indicating higher pigmentation (McGraw et al., 2005). The spectral curves of the head, flank and chin of males and the chin and cheek of females respectively, revealed the presence of plumage components facilitating the reflectance in the UV-range (Figure 2). In order to quantify UV-reflectance of the feathers, we calculated UV-Chroma, which is the proportion of reflectance in the UV-range (320–400 nm) compared to the total reflectance (320–700 nm) (Griggio et al., 2010a, Hill & McGraw, 2006b, Mahr et al., 2012). Spectral measurements were restricted to a range between 320-700 nm, which reflects the avian color vision spectrum (Hill and McGraw 2006).

In addition to the colorimetric variables, we measured the tail length for both sexes and quantified the length of the black male beard, which were previously shown to be sexually selected traits in this species (Hoi & Griggio, 2008). Tail length was measured using a ruler (to the nearest 0.5 mm). To measure the beard length (to the nearest 0.01mm), three photographs (left/right side and front) of each male individual were taken by placing the birds in front of a millimeter paper at a standardized distance (25 cm) from the camera (Nikon D 60, Nikon Corporation, Tokyo, Japan) mounted on a tripod (Manfrotto, Vitec imaging solutions spa, Cassola, Italy). We used the software ImageJ (Rueden et al., 2017) to determine the beard length, by measuring a known distance on the millimeter paper as a scale (10 mm on each picture) and calculating the distance between the lowest and highest point of the beard in mm. For the analyses we calculated the average beard length (in mm).

### IGF-1 plasma concentrations

Plasma IGF-1 levels were measured in duplicates by a competitive ELISA developed in our laboratory at University of Debrecen (Lendvai et al. unpublished data). An assay kit is available upon request from the authors. 96-wells NUNC microplates were coated overnight at 4°C with 100μl of an antibody raised against IGF-1 in rabbits. The capture antibody was incubated for 2 hours at room temperature (24°C) with 20 μl known concentrations (in serial dilutions starting at 500 ng/ml) of synthetic chicken IGF-1 or 20μl of sample and 100 μl biotinylated IGF-1 as a tracer. After incubation, the microplate was washed three times with 250 μl of PBS buffer (8 g NaCl, 0.2 g KCl, 1.44 g Na_2_HPO_4_ and 0.24 g KH_2_PO_4_ in 1000 ml ddH2O, pH 7.4) containing 0.025% Tween 20. After washing, 100 μl of streptavidin-horseradish peroxidase conjugate was added to all wells and incubated at room temperature at 30 mins, followed by another washing cycle (3 times). Then, 100 μl of tetra-methyl-benzidine was added to the wells and incubated at room temperature for 30 minutes. The enzymatic reaction was stopped by adding 100 μl of 1M H_2_SO_4_, and optical density was measured at 450 nm (reference at 620 nm). The calibration curve was fitted using a 4-parametric log-logistic curve, and concentrations of unknown samples were read off from this curve. We used a chicken plasma in quadruplicate to determine intra- and inter-assay coefficient of variation (4.8% and 9.7% respectively). In this assay, we did not use any extraction protocol on our samples, because we were interested in the free, biologically active fraction of IGF-1, which is not bound to IGF-1 binding proteins.

### Statistical analysis

In total n = 23 males and n = 20 females were sampled. Statistical analyses were conducted using Statistica 7.1 (Statsoft Inc., Tulsa). However, we excluded one male from the analyses of the plumage coloration, because we did not achieve a sufficient number of measurements.

To test the relationship between the different ornamental features in males and females, we conducted principal component analyses (PCAs) for each sex separately. Therefore we combined male tail length, beard length and the spectral variables (Brightness and UV-Chroma), describing the coloration of the head, chin, flank and back into a PCA. We applied the same procedure to females using tail length and colorimetric variables describing the coloration of the female cheek, chin and back. We applied ‘varimax’ rotations and normalized the data. The factor scores of the consecutive PCs explaining the highest variation within the chosen variables, were extracted using the Kaiser criterion (eigenvalues higher than 1). Based on the findings of the PCA (see results section), we conducted a second PCA on the UV-Chroma variables of different body parts to produce a single variable representing the structural plumage coloration (Males: eigenvalue = 2.62, variance explained = 87.46%; Females: eigenvalue = 1.51, variance explained = 75.28%) for further analyses. We refer to this variable as “overall UV-Chroma”. We used GLMs to explore the relationship between baseline IGF-1 levels, structural components of the plumage coloration (overall UV-Chroma), brightness, beard length and the morphological plumage trait (tail length). Each of these variables were tested separately with each plumage trait entering the initial model as dependent variable and IGF-1 being the explanatory variable. Tarsus length and body-mass were incorporated as covariates. Since IGF-1 might be linked to the nutritional state of individuals, we also included an interaction between body-mass and IGF-1 levels.

Model selection was conducted using stepwise backward method. Starting with the interaction, non-significant terms were eliminated step-by-step from the model. Only significant variables were retained in the final model and each removed variable was re-entered separately into the final model to test their effects (Engqvist, 2005, Grafen & Hails, 2002, Mahr et al., 2016). We provide parameter estimates *±* SE and two-tailed tests throughout.

## Results

### Integration of multiple plumage ornaments

PC1 and PC2 captures most variation in plumage coloration of the colorimetric variables (PC1: Eigenvalue = 3.39, total variance = 56.57%; PC2: Eigenvalue = 1.13, total variance = 18.86%). Brightness of the back loads negatively, whereas UV-Chroma of the same areas loads strongly positively to PC1, suggesting that high reflectance of structurally based components of the plumage indicate low color intensity (Table 1). The three UV-Chroma variables show very similar loadings. PC2 explains most variation in the beard length (Table 1), whereas tail length loadings differs from both colorimetric variables and beard length. Considering these factor loadings there is strong indication that plumage coloration, ornamental patterns (beard length) and tail length are independent traits (Table 1).

**Table 1:**
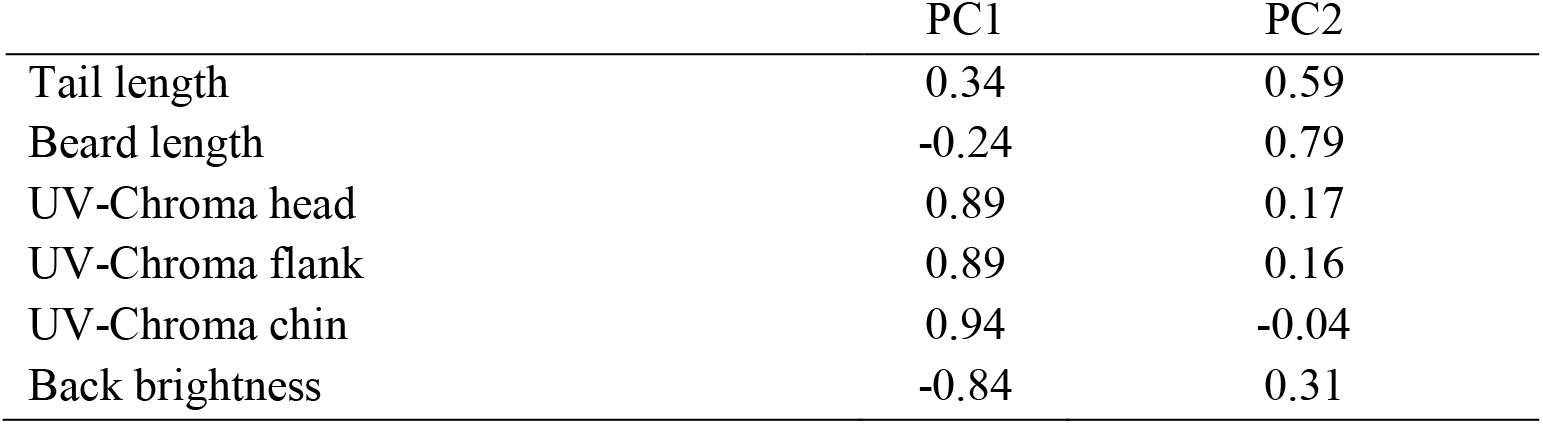
Loadings of PC1 and PC2 and in male bearded reedlings

|  | PC1 | PC2 |
| --- | --- | --- |
| Tail length | 0.34 | 0.59 |
| Beard length | -0.24 | 0.79 |
| UV-Chroma head | 0.89 | 0.17 |
| UV-Chroma flank | 0.89 | 0.16 |
| UV-Chroma chin | 0.94 | -0.04 |
| Back brightness | -0.84 | 0.31 |

Female coloration showed a similar pattern to males: PC1 (Eigenvalue = 1.57, total variance = 39.3%) and PC2 (Eigenvalue = 1.18, total variance = 29.45%). Similarly to males, the two UV-Chroma variables show very similar loadings, which is perpendicular to the brightness of the back and opposite from tail length, indicating that brightness of the back and UV-Chroma of the structural plumage components are independent traits, whereas tail length is negatively related to color intensity in females (Table 2).

**Table 2:** Loadings of PC1 and PC2 in female bearded reedlings

|  | PC1 | PC2 |
| --- | --- | --- |
| Tail length | -0.15 | 0.76 |
| UV-Chroma chin | 0.85 | -0.02 |
| UV-Chroma cheek | 0.87 | -0.07 |
| Brightness back | -0.07 | -0.80 |

### Do IGF-1 levels predict plumage colors, tail length and ornamental patterns?

Male tail length was positively related to IGF-1 (*F*_1,21_ = 6.65, *β* = 0.49 *±* 0.19, *p* = 0.02, Figure 3a). In contrast, female tail length was not affected by IGF-1 levels (*F*_1,17_ = 0.23, *p* = 0.64). However, female tail length tended to be positively related to body mass during molt (*F*_1,17_ = 4.07, *p* = 0.06).

**Figure 3.**
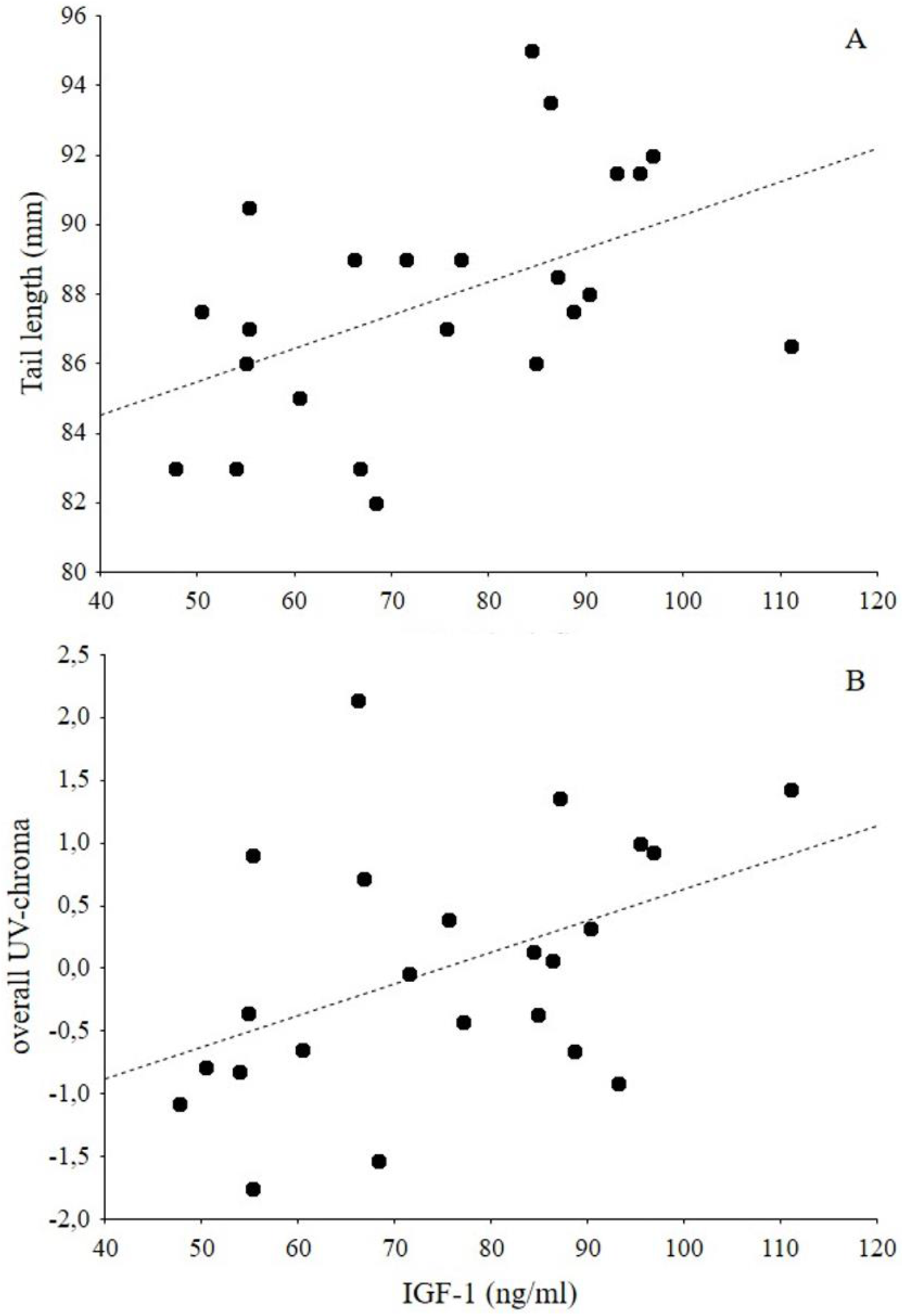
Baseline IGF-1 levels during molt are related to sexually selected ornaments in male bearded reedlings. Males with higher IGF-1 levels have (a) longer tails and (b) develop plumage with higher reflectance in the UV-range.

Beard length was not related to any of the variables measured during the molt (tarsus length: *F*_1,20_ = 0.51, *p*= 0.48, body mass: *F*_1,20_ = 1.36, *p* = 0.26, IGF-1 levels: *F*_1,20_ = 0.26, *p* = 0.62). The brightness of the male back plumage was also unrelated to tarsus length (*F*_1,21_ = 0.003, *p* = 0.95), body mass (*F*_1,21_ =0.79, *p*= 0.39) or IGF-1 levels (*F*_1,21_ = 1.53, *p* = 0.23). However, we found a positive relationship between IGF-1 levels and overall UV-Chroma in males (*F*_1,21_ = 5.09, *β* = 0.44 *±* 0.19, *p* = 0.03, Figure 3b).

In females, the brightness of the back showed a complex relationship with body mass and IGF-1 levels. Heavy birds with low IGF-1 levels and light birds with high IGF-1 levels had darker back coloration than heavy birds with high IGF-1 or light birds with low IGF-1, indicated by a significant effect of body mass (*F*_1,14_ = 8.90, *β* = − 2.89 *±* 0.97, *p* < 0.01), IGF-1 (*F*_1,14_ = 9.27, *β* = − 10.67 *±* 3.5, *p* < 0.01) and their interaction (*F*_1,14_ = 8.84, *β* = 10.06 *±* 3.39, *p* = 0.01).

Overall female UV-Chroma was not related to either IGF-1 (*F*_1,16_ = 0.532, *p* = 0.48), body mass (*F*_1,16_ = 2.84, *p* = 0.11) or the interaction term IGF-1*body mass (*F*_1,16_ = 1.26, *p* = 0.28). Tarsus length was also unrelated to the UV plumage components in females (*F*_1,16_ = 1.78, *p* = 0.20).

## Discussion

This study provides the first evidence that IGF-1 may facilitate the development of sexually selected plumage traits in birds. Our results indicate that in male bearded reedlings IGF-1 levels during molting predict tail length and UV-reflectance of the structural components of plumage (Figure 3). We further demonstrate that IGF-1 affects multiple ornamental plumage traits differently in the two sexes. Our findings are corroborated by previous studies on bearded reedlings and confirm that different ornamental features, such as plumage colors and tail length do not correlate. This raises the idea that they either reveal a different set of information (rather than amplifying another signal), or carry no further informational content (Griggio et al., 2016, Hoi & Griggio, 2008). Within this context, the question arises how IGF-1 can affect the development of a sexually selected character and whether it provides a causal link between body-condition and parameters of plumage quality.

Molting is an energy demanding process, with concomitant major metabolic changes and an increased demand for protein synthesis, cell growth, proliferation and differentiation (Kuenzel, 2003). The corresponding physiological requirements might be an important factor, linking feather development to parameters of individual condition before and during the molt (Jovani & Blas, 2004, Murphy et al., 1988, Pap et al., 2008). In white crowned sparrows (*Zonotrichia leucophrys gambelii*) and house sparrows (*Passer domesticus*) for example, malnutrition had severe effects on the quality of feathers and the length of flight feathers (Murphy et al., 1988, Pap et al., 2008). In addition, late hatching, late arrival from overwintering sites and poor body condition, often force individuals to increase molt speed to keep pace with conspecifics and to be able to compete for mating partners or resources. This compensatory measure has negative effects on feather length and plumage quality (Dawson et al., 2000, Vágási et al., 2012). IGF-1 is highly responsive towards the availability of environmental resources, due to its sensitivity to the nutritional status of an individual and to dietary components (e.g. proteins). It can affect whether energy is allocated into cell proliferation, growth and protein syntheses (Blumenthal et al., 2011, Dantzer & Swanson, 2012, Tighe et al., 2016), which might constitute an important adaptive adjustment to the environmental conditions with long lasting effects on individual life-history (Dantzer & Swanson, 2012, Holzenberger et al., 2003, Lewin et al., 2017). The same properties might not only affect growth and life span, as was shown previously (Dantzer & Swanson, 2012), but might also have the potential to regulate the allocation of energy into feather growth. Hence, IGF-1 might provide a potential link between resource availability, body condition and plumage traits. Higher levels of IGF-1 during the molt might stimulate feather development and facilitate the display of more elaborated plumage features. Indeed, we show that in bearded reedlings, IGF-1 correlates with male, but not female tail feather length and UV-reflectance. The development of structural plumage components is complex and requires the regulation of protein synthesis and breakdown of keratin (Hill & McGraw, 2006a, Hudson & Wilcoxen, 2018). There is strong indication that, similarly to longitudinal growth of feathers, the growth of the regularity of the nanoscaled structures, which cause the reflection in the UV-range is linked to parameters of individual physiological condition during the molt. This was shown in different species, for instance, in blue tits (*Cyanistes caeruleus*), accelerated molt causes a decrease in the saturation of the UV-blue crown feathers (Griggio et al. 2009). Also in dark eyed juncos (*Junco hyemalis*) the availability of resources (dietary restrictions) during molting negatively affected the growth of structural plumage components (McGlothlin et al., 2007).

In brown-headed cowbirds (*Molothrus ater*), structural plumage traits, but not melanin based plumage coloration were shown to be affected by nutritional stress (McGraw et al., 2002).

These findings support our results, which did not reveal a relationship between the measured melanin based ornamental features (beard length and brightness of the back), IGF-1 levels, body mass or tarsus length. This is particularly interesting because the beard of male bearded reedlings is a melanin-based plumage ornament, which underlies strong sexual selection processes. Males with longer beards are more dominant and females clearly display a preference for them as potential mates (Griggio et al., 2016, Hoi & Griggio, 2008, Hoi & Griggio, 2012). These characteristics raise the idea that the beard might be costly to produce and serves as honest indicator of quality (Andersson, 1994b, Griggio et al., 2016, Hoi & Griggio, 2008). Indeed, the degree of melanization and the size of some types of melanin-based ornaments can be sensitive towards environmental factors, fluctuating testosterone levels and dietary components and hence serve as potential indicator for condition during the molt (Jawor & Breitwisch, 2003, Musgrove & Wiebe, 2016, Roulin, 2004, Roulin, 2016). However, it was previously suggested that some types of melanin-based ornaments are considered as relatively cheap to produce and they underlie other physiological constrains and genetic prerequisites (Roulin, 2016, Senar et al., 2003). The different physiological properties of the measured ornamental plumage traits might therefore explain that structural, but not melanin-based ornaments were affected by circulating IGF-1 levels in bearded reedlings.

In females, neither of the ornamental features were significantly correlated with IGF-1 levels. This is particularly surprising, because tail length underlies mutual mate choice in bearded reedlings and we found a weak correlation between tail length and body mass, which corresponds with the idea that it serves as potential indicator for quality. Interestingly, our study indicates that dependent on the body-mass, IGF-1 might positively affect the melanin pigmentation of the brown back in females, whereas no such relationship became apparent in males. Heavier females with lower IGF-1 and smaller females with high IGF-1 displayed darker back-plumage than heavy birds with high IGF-1 or light birds with low IGF-1. One possible explanation is that this effect is due to different selection pressure acting on males and females (Romero-Pujante et al., 2002). We cannot rule out that IGF-1 might have effects, which facilitate the growth of more pigmented feathers in females, but it should be considered that overall, our study did not reveal a significant relationship between body mass or size during the molt and any of the plumage ornaments in both sexes. However, we obtained only one measure of body-condition and molting birds undergo an energetic demanding period, which was previously demonstrated to significantly affect parameters of body condition. Muscle score, body fat and mass in molting birds are often low (Dolnik & Valery, 1979, Minias et al., 2010, Swaddle & Witter, 1997) which may explain why we did not find a relationship between body mass and the ornamental features.

Overall, our study is the first to report a relationship between IGF-1 levels during molting and the elaboration of ornamental feather traits and colors in adult birds. The distinct properties of IGF-1, namely its sensitivity towards nutrition and its important role in the mechanisms stimulating growth and development, make it a potential candidate to further investigate the mechanisms linking parameters of individual condition and the capacity to display elaborated ornaments.

## Acknowledgments

We thank S. Wiedermann and A. Nurmisto for their support during the data collection and G. Rédai for painting Fig. 1. We further thank the Nationalpark Neusiedlersee for access to the study population and hosting our project during the fieldwork. Funding was provided by the Hungarian National Development, Research and Innovation Office (NKFIH) (OTKA K113108) and by a bilateral research grant of the NKFIH (TÉT 15-1-2016-0044) and the Österreichischer Austauschdienst (ÖAD, WTZ #HU 02/2016). KM was supported by a Schrödinger fellowship (# J4235-B29) granted by the FWF. AZL and ZT were supported by a grant from the European Union and the European Social Fund (EFOP-3.6.1-16-2016-00022) and AZL by the Romanian Ministry of Education (PN-III-P4-ID-PCE-2016-0572).

